# *De novo* SNP calling reveals the candidate genes regulating days to flowering through interspecies GWAS of *Amaranthus* genus

**DOI:** 10.1101/2021.10.05.463269

**Authors:** Ya-Ping Lin, Tien-Hor Wu, Yan-Kuang Chan, Maarten van Zonneveld, Roland Schafleitner

**Author notes:** **Correspondence:** Corresponding author: Ya-Ping Lin, Roland Schafleitner.

## Abstract

Amaranths serve as pseudo cereals and also as traditional leafy vegetables worldwide. In addition to high vigor and richness in nutrients, drought and salinity tolerance of amaranth makes it a promising vegetable to acclimatize to the effects of global climate change. The World Vegetable Center genebank conserves about 1,000 amaranth accessions and various agronomic properties of these accessions were recorded during seed regeneration over decades. In this study, we verified the taxonomic annotation of the germplasm based on a 15K SNP set. Besides, in the assumption that the yield components of grain amaranths are different from those of leaf amaranths, we observed that grain amaranths presented larger inflorescences and earlier flowering than leaf amaranths. Dual-purpose amaranth showed larger leaves than leaf amaranths and later flowering than grain amaranths, which seemed reasonable because farmers can harvest more leaves during the prolonged vegetable stage, which also provides recovery time to enrich grain production. Considering frequent interspecies hybridization among grain amaranth complex, we proposed an interspecies GWAS for days to flowering, identifying a *AGL20/SOC1* homolog. Meanwhile, another GWAS using only *A. tricolor* accessions revealed six candidate genes homologous to *lba1*, *bri1*, *sgs1* and *fca*. These homologous genes were involved in the regulation of flowering time in Arabidopsis. This study revealed the usefulness of genotypes for species demarcation in the genus *Amaranthus* and the potential of interspecies GWAS to detect QTLs across different species, opening up the possibility of targeted introduction of specific genetic variants into different *Amaranthus* species.

## Introduction

The genus Amaranthus comprises about 50 species, most of them annual weeds known as pigweed, that are distributed worldwide (Bonasora et al., 2013). Most likely, amaranths were already collected by humans as a food source for a very long time before some species have been domesticated for grain production by Aztec and Mayan civilization (Lucian et al., 2018; Marx, 1977; Sauer, 1950, 1967; Tucker, 1986). In Asia, amaranths were domesticated as a leafy vegetable. Today, a complex of *A. caudatus, A. cruentus* and *A. hypochondriacus*, serve as pseudo cereals in America and Africa, and *A. cruentus* have evolved into leaf amaranth after it was introduced to Africa; meanwhile, *A. blitum*, *A. dubius*, *A. tricolor* and *A. viridis* are cultivated or collected from the wild as traditional leaf vegetables in Africa and Asia (Das 2016, van Zonneveld et al., 2021). The amaranth crop has the potential to convey a range of benefits without high inputs (Joshi et al., 2018; Tucker, 1986). It can be cultivated year-round with short rotations and with low input even under harsh conditions. As a C4 plant, it can prevent water loss during photosynthesis, serving as a resilient drought and salinity-tolerant crop with potential to adapt well to some climate change scenarios (Huerta-Ocampo et al., 2009, 2011; Joaquín-Ramos et al., 2014; Sarker et al., 2020; Sarker and Oba, 2020; Wang et al., 1992, 1993). In terms of nutrients, amaranth grain is rich in proteins with balanced amino acid content, minerals and dietary fibers, and it is free of gluten, providing a dietary option for gluten-intolerant consumers; meanwhile, amaranth leaves are rich in vitamins and minerals, including Ca, Fe and Zn (Coelho et al., 2018; Miranda-Ramos et al., 2019; Prakash and Pal, 1991; Valcárcel-Yamani and Lannes, 2012; Venskutonis and Kraujalis, 2013). Oxalate content may reduce mineral uptake, but adapted cooking methods can prevent this effect (Yadav and Sehgal, 2003). Being an orphan crop, genetic research of *Amaranthus* species has been neglected, narrowing the scope of amaranth conservation and utilization.

The genome size of *Amaranthus* species ranges from 500 Mb to 800 Mb (Stetter and Schmid, 2017), and the basic chromosome set of diploid amaranth species is 2n = 32 or 2n =34 (Madhusoodanan and Pal, 1981; Greizerstein et al., 1997; Greizerstein and Poggio, 1994; Bonasora et al., 2013). *A. dubius*, a tetraploid amaranth species, has a chromosome number of 2n = 64 (Grant, 1959). Despite different chromosome numbers among the diploid *Amaranthus* species, the possibility of interspecies hybridization has revealed promising heterosis in hybrids. For example, high-parent heterosis of up to 40% was found for biomass in the progeny derived from *A. hypochondriacus* x *A. cruentus* (Lehmann et al., 1991). *A. hybridus* was suggested to be the ancestor of the grain amaranth complex and it was possible to cross this species with *A. caudatus, A. cruentus* and *A. hypochondriacus* (Kietlinski et al., 2014; Stetter et al., 2016). The progeny derived from parents with different chromosome number had 2n = 32, 33, 34 in a ratio of 1 : 2 : 1, implying the potential of *A. hybridus* as a bridge crossing species (Pal et al., 1982), which may be useful for breeding interspecific hybrids.

One of the consequences of little systematic investigation in the *Amaranthus* genus is its confusing species classification. Plant taxonomists identified *Amaranthus* species mainly according to a few floral characteristics (Sauer, 1950, 1967). In addition, interspecies hybridization and plant plasticity make the morphological-based classification ambiguous (Achigan-Dako et al., 2014; Pal and Khoshoo, 1972, 1973; Pal et al., 1982; Stetter and Schmid, 2017). Based on molecular markers, such as isozymes or low-copy genes, the phylogenetic relationship among the species of the *Amaranthus* genus was not conclusive, implying the demand of more markers for classification (Chan and Sun, 1997; Waselkov et al., 2018). Recently, Stetter and Schmid (2017) used genotyping-by-sequencing to obtain 3K SNPs for 35 *Amaranthus* species and constructed a species tree, illustrating that the grain amaranth complex belonged to one clade, while leaf amaranth species *A. tricolor*, *A. blitum*, and *A. viridis*, as well as *A. dubius* were separated into different clusters, and the latter was close to the grain amaranth complex. Besides, this study also indicated that the reference genome-based SNP calling for genotyping many species results in a loss of SNPs compared to *de novo* SNP calling, because genome size, chromosome number and likely also sequence varied among these *Amaranthus* species. When SNP calling for various *Amaranthus* species was based on the reference genome of *A. hypochondriacus* (Lightfoot et al., 2017), which comprises about 500 Mb, many reads remained un-mapped on the reference genome, resulting in fewer SNPs than *de novo* SNP calling. This result suggested that for genotyping multiple amaranth species, *de novo* SNP calling is preferable to obtain a universal SNP dataset, which increases the SNP number and therefore benefits the mining of biodiverse genetic resources. Meanwhile, only a few studies investigated leaf amaranths systematically (Nguyen et al., 2019). Considering the richness in nutrients, the resilience to climate stress and the overall potential to harness food production for changing climates, amaranth deserves more attention.

The conservation of germplasm is the core mission of genebanks. Over decades of germplasm collection, regeneration and characterization, the collection sites and taxonomic annotation were documented and morphological descriptors were recorded and stored as passport data information for genebank accessions. However, due to limited resources, especially for orphan crops, data collection can be limited. Species attribution often was restricted to genus than to the species level. Genetic analysis, including genome-wide association study (GWAS) in genebank collections of orphan crops is often constrained by the small size of collections for a single species, which reduces the detection power for rare alleles (Korte and Farlow, 2013). To increase the frequency of minor alleles, several close species could be merged together in a single GWAS, but this may introduce population structure and result in confounding effects and a high false discovery rate (Huang and Han, 2014). Investigation of a natural population generates a general dilemma: if emphasis is given to locally relevant samples without population structure, such as a single species, detection power remains low due to small population size. If a broader sample is chosen, population size and detection power increases, but population structure is introduced (Brachi et al., 2011). Progress in developing models for correcting population structure have yielded promising results to identify quantitative trait loci in interspecific diversity panels, indicating that the confounding effect through population structure can be overcome in GWAS (Bauchet et al., 2017; Noble et al., 2018; Zhao et al., 2019). Under the concept of comparative genomics, individuals of close species can be resequenced and their sequences can be mapped back to the same reference genome or used for *de novo* SNP calling to investigate traits in a large diverse panel.

So far, most genebanks mainly hold grain amaranth species, especially *A. hypochondriacus* (Genesys, 2017; Joshi et al., 2018). As one of the largest public sector vegetable genebanks, the World Vegetable Center maintains nearly 1,000 *Amaranthus* accessions, and more than half of them are classified as vegetable (leafy) amaranths. This collection provided us an opportunity to investigate amaranth as a vegetable and dual-purpose crop that has a completely different harvest model than grain amaranth (Dinssa et al., 2018; Hoidal et al., 2019). In this study, we used *PstI-MseI* double-digest Restriction site-Associated DNA sequencing (ddRAD sequencing) to perform *de novo* SNP calling. A universal SNP dataset across *Amaranthus* species was obtained. The universal SNP dataset allowed us to revisit the taxonomic annotation of *Amaranthus* accessions and to investigate the genetic diversity of the major species in the collection of the World Vegetable Center. In addition, candidate loci that regulate days to flowering were revealed by interspecies GWAS.

## Materials and Methods

### Plant materials and morphology records

The World Vegetable Center genebank collection contains 13 *Amaranthus* species, but most accessions belong to eight species: *A. blitum*, *A. caudatus*, *A. cruentus*, *A. dubius*, *A. hypochondriacus*, *A. spinosus*, *A. tricolor* and *A. viridis* (Figure 1). The genebank accessions available at https://avrdc.org/seed/unimproved-germplasm/ were characterized during seed regeneration for several agronomic traits from 1995 to 2019 in Shanhua, Taiwan. The collected data were used in this study (Table S1). All the traits were documented following the Amaranthus morphological descriptors (http://seed.worldveg.org/public/download/descriptors/Amaranthus_2015.pdf.). To control for uncertainty and bias due to variable climatic conditions and changes in crop management and the lack of control accessions carried over the years, we performed Principal Component Analysis (PCA) of both the agronomical traits (Table S1) and the climatic data from a weather station located in proximity of the regeneration plots, including mean, maximum, minimum temperature, precipitation, evaporation, mean humidity, mean soil temperature under 10 cm and 30 cm, and solar intensity (Table S2). After ensuring that there was no obvious phenotypic or climatic clustering over the years, we combined all phenotyping data recorded over decades together in the following analyses. PCA was performed to observe the variation of the important morphological data among different *Amaranthus* species, including plant height (descriptor identifier Am220), mean length of basal lateral branches (Am240), mean length of top lateral branches (Am250), leaf length (Am290), leaf width (Am300), days to flowering (Am400), terminal inflorescence stalk length (Am410), terminal inflorescence laterals length (Am420), length of axillary inflorescence (Am460), seed shattering (Am500) and 1000 seed weight (Am540; Table S1). PCAs were completed by R package “factoextra” (Kassambara and Mundt, 2020).

**Figure 1.**
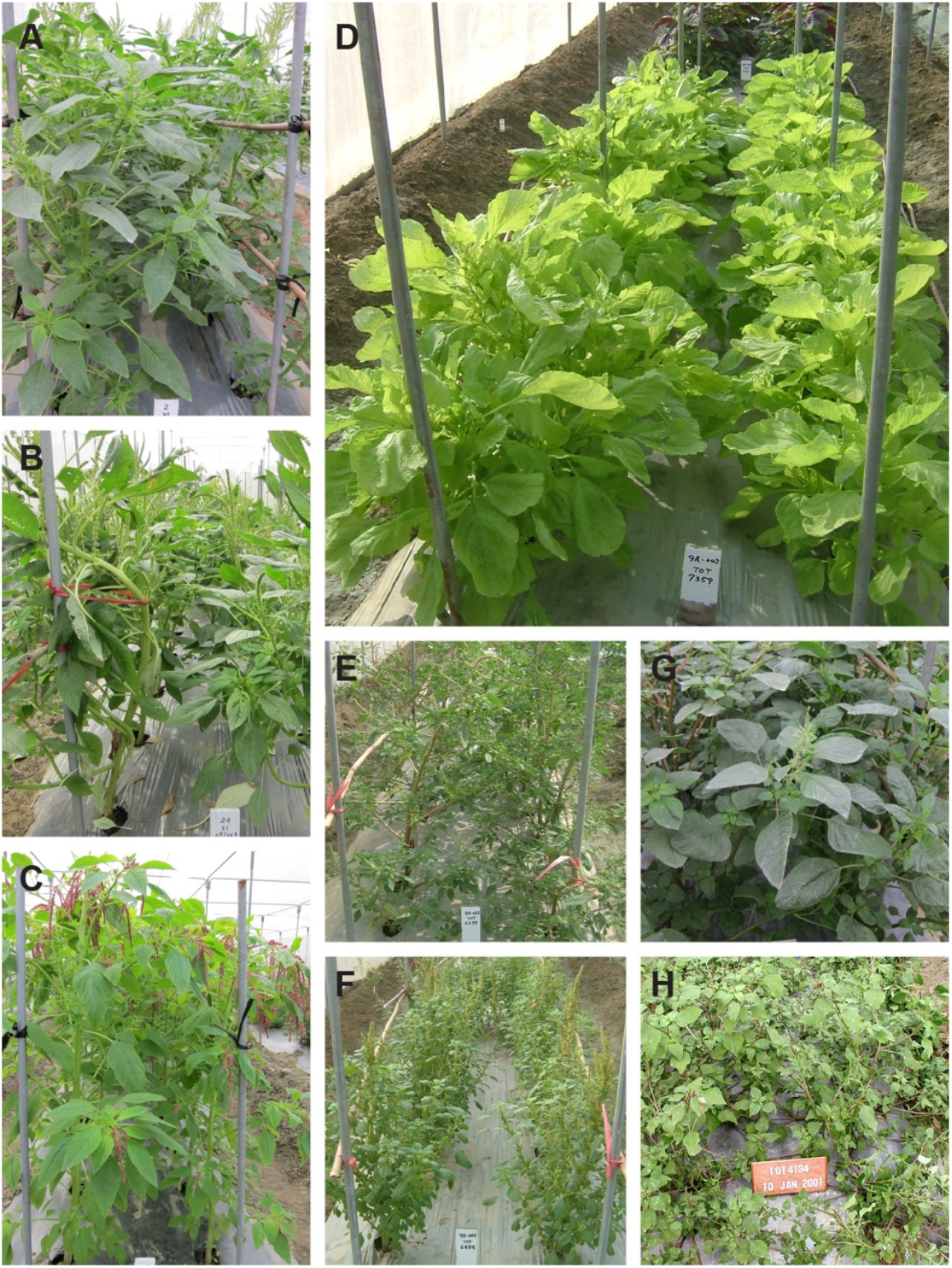
Representative plant images of eight major *Amaranthus* species of the World Vegetable Center germplasm collection. (A) *A. cruentus* (B) *A. hypochondriacus* (C) *A. caudatus* (D) *A. tricolor* (E) *A. spinosus* (F) *A. blitum* (G) *A. dubius* (H) *A. viridis*.

### DNA Sequencing

In a total of 731 accessions out of 777 accessions available in 2017 (current collection size: 1,062 accessions) germinated and were used for ddRAD sequencing. For each accession, freeze-dried young leaf tissue of a single plant was collected and genomic DNA was extracted with sbeadex maxi plant kit (LGC, UK) according to the supplier manual. DNA integrity and quality were evaluated by electrophoresis on 1% agarose gels and DNA was quantified on a fluorimeter (Qubit). The samples were sequenced by Diversity Arrays Technology Pty Ltd. (DArT, Australia) using DArTSeq™. DArTseq technology provides a selective genome sequencing method by an optimized combination of restriction enzyme digestion, which in our case was *PstI*-*Mse*1. All the sequences are available in the NCBI BioProject PRJNA734966.

### SNP calling

After sequencing reads filtering by SolexaQA with Phred quality score threshold of ≥ 20 (Cox et al., 2010), we performed a *de novo* SNP calling using the Stacks software version 2.54 (Catchen et al., 2013). The reason we applied *de novo* SNP calling was because we intended to obtain a universal SNP set across several different *Amaranthus* species in this study and for most of these species no reference genome was available. If sequencing reads derived from various species would have been mapped to a reference genome of a single species, the portion of reads derived to species-specific genomic segments would have been lost for SNP calling (Stetter and Schmid, 2017). In addition, according to the principle of comparative genomics, common characteristics of related species are often controlled by conserved sequences. These conserved sequences could be assembled into the loci matching to the same read catalogs through *de novo* assembly using the “denovo-map.pl” pipeline in Stacks. The *de novo* SNP calling approach was applied with a missing data rate threshold of < 0.5 and a heterozygosity proportion of < mean + 3*standard deviation. The resulting universal SNP dataset contained a total of 15,573 SNPs. We also performed reference genome-based mapping back on the *A. hypochondriacus* reference genome by bwa (Li and Durbin, 2009; Lightfoot et al., 2017). The percentage of mapped reads among different species was compared between these two approaches.

### Genetic structure

To investigate the genetic structure in the collection, we first performed a PCA using the universal SNP dataset based on genetic distance calculated in TASSEL 5.0 (Bradbury et al., 2007). A phylogenetic tree was constructed in TASSEL 5.0 and plotted using R package “phytools” (Revell, 2012). Structure analysis for K values from 2 to 20 was completed with 10 replications in R package “LEA” (Frichot and François, 2015). According to the cross-validation, we used the smallest K that can separate grain amaranth species as the best K.

### Correction of the taxonomic annotation of the amaranth germplasm

The *Amaranthus* taxonomy can be very complex and confusing due to the few characteristics that distinguish species, frequent interspecies hybridization and phenotypic plasticity (Pal et al., 1982; Pal and Khoshoo, 1972, 1973; Sauer, 1950, 1967; Stetter and Schmid, 2017). Therefore, after the best K was determined, we used the genetic component matrix (Q matrix) as a reference to re-examine the species annotation in the passport data set of the amaranth collection. We first used Q component > 0.9 as a threshold to summarize the major species for each Q component, resulting in one Q component corresponding to one specific species. Afterwards, we re-evaluated the plant images of these accessions that showed inconsistency in their taxonomy between the passport data and the Q component. The annotation was only corrected in accessions with a Q component > 0.9 and that could be re-classified based on the plant images. For the accessions with Q components < 0.9, it was preferred to treat them as intermediate types and not to correct their taxonomic annotation in the passport data.

### Genetic diversity

After determination of the genetic groups, we used the accessions that had a single Q component > 0.9 and were consistently classified based on genetic components and morphological descriptor to a specific taxonomic group to investigate the genetic diversity. The analysis of molecular variance (AMOVA) was estimated by the R package “hierfstat” to quantify the population differentiation (Goudet, 2005). Besides, the observed heterozygosity (Ho) and the gene diversity within-population (Hs) were calculated by the R packages “adegenet” (Jombart and Ahmed, 2011). The number of private alleles was calculated by the R packages “poppr” (Kamvar et al.,2015). segregation site (S), nucleotide diversity (π) and calculated nucleotide diversity (θ) were estimated with TASSEL 5.0 (Bradbury et al., 2007).

### GWAS of days to flowering

GWAS among several related species (interspecies GWAS) is a common strategy to identify QTLs that contribute to important agronomic traits in crops. This strategy could not only include more diversity into the genetic and phenotypic analyses, but also increases the sample size of target populations. Following the same logic, interspecies GWAS was performed here to overcome the complicated taxonomy in the genus Amaranthus and to compensate for the small collection size of most individual *Amaranthus* species. We used the filtered SNPs with minor allele frequency < 0.05 to perform GWAS for two diverse panels, the grain amaranth complex and *A. tricolor*, in TASSEL 5.0 (Bradbury et al., 2007). Four models, including Phenotype = Genotype (without any correction), Phenotype = Genotype + PC1 matrix, Phenotype = Genotype + Q matrix and Phenotype = Genotype + Kinship, were completed. Models without kinship were general linear models (GLM). The trait was transformed with log10 before proceeding with GWAS. The Q-Q plots were plotted by R package “qqman” (Turner 2017). The false discovery rate (FDR) was calculated by R function “p.adjust” (Benjamini and Hochberg). A significant SNP was defined with FDR < 0.05.

### Screen for candidate genes contributing to days to flowering

Given that days to flowering is a well-studied trait in plants, we used this trait for the further candidate gene survey in the interspecies GWAS. We extracted the sequence catalog of the significant SNPs responding to days to flowering and blasted them on the reference genome of *A. hypochondriacus* (Lightfoot et al., 2017). As *de novo* SNP calling could not estimate the decay of linkage disequilibrium (LD), we followed the LD decay estimations for the whole genome sequence of Arabidopsis (Kim et al., 2007) and used 10-kb segments around the significant SNPs to screen for candidate genes in the presumable LD intervals. Based on the genome annotation of *A. hypochondriacus*, genes located from 10-kb upstream to 10-kb downstream of the significant SNPs were blasted in a threshold of *e* value < 10e-5 in the Arabidopsis protein database (TAIR10). All the blast works were completed with “ncbi-blast-2.11.0” (Camacho et al., 2009). We screened the annotations of these homologous genes manually to identify the candidate genes participating in the regulation of flowering time.

## Results

### Comparison of SNP calling by *de novo* assembly with reference genome-based SNP calling

The average reads obtained per genotype were 1.2 million, ranging from 1 million to 1.5 million per accession (Table S3). In average, the percentage of mapped reads in a reference genome-based and *de novo* assembly was 67.41% and 74.60%, respectively (Table S3). The percentage of reference genome-based mapping of the *A. hypochondriacus* accessions was 80.37% in average, while that of the rest of the accessions was 66.96%. In comparison, the percentage of reads assembled *de novo* assembly for the *A. hypochondriacus* accessions was 76.91% and that of the rest of the accessions was 74.32% (Table 1 and Table S3). *A. tricolor* makes up the largest part of the WorldVeg Amaranthus collection, and only 59.75% of the sequencing reads of accession of this species accessions could be mapped to the *A. hypochondriacus* reference genome, while 73.24% of the reads could be mapped in the *de novo* assembly (Table 1). This suggested that *de novo* assembly resulted in more complete assembly of reads than reference genome-based mapping, especially for accessions of species that are more divergent from the species of the reference genome. Therefore, we used *de novo* assembly with an average coverage of 27-fold to develop the SNP markers for this study. The missing data rate ranged from 0.11 to 0.41 and the heterozygosity ranged from 0.00 to 0.31 (Table S3).

**Table 1.** The summary of reads mapped by reference-based and *de novo* assembly approaches

| Species | Reference-based<br>mapped reads | Reference-based<br>mapped percentage | De novo assembly<br>reads | De novo assembly<br>percentage |
| --- | --- | --- | --- | --- |
| *A. blitum* | 878,005 | 76.02 | 886,808 | 76.58 |
| *A. caudatus* | 1,014,530 | 78.13 | 993,968 | 76.48 |
| *A. cruentus* | 1,030,248 | 79.88 | 968,266 | 75.10 |
| *A. dubius* | 897,324 | 71.98 | 972,506 | 78.02 |
| *A. hybridus* | 745,556 | 71.15 | 784,840 | 74.90 |
| *A. hypochondriacus* | 1,035,106 | 80.37 | 990,408 | 76.91 |
| *A. palmeri* | 769,962 | 55.97 | 1,027,511 | 74.70 |
| *A. spinosus* | 911,194 | 69.19 | 933,853 | 70.86 |
| *A. thunbergii* | 864,583 | 67.56 | 1,035,079 | 80.67 |
| *A. tricolor* | 781,028 | 59.75 | 956,670 | 73.24 |
| *A. viridis* | 827,814 | 62.51 | 968,842 | 73.18 |

### Genetic groups of *Amaranthus* genus

The taxonomy in the genus *Amaranthus* may cause difficulties for genebank curators due to very few key distinct traits among the species (Sauer, 1950, 1967). Here, we used genetic clusters to verify the species information in the passport data of the *Amaranthus* genebank accessions. According to the PCA of the eight major species in the WorldVeg collection (Figure 2A), grain amaranth species were genetically close to each other, while vegetable amaranth species were separated into distinct clusters. In addition, *A. cruentus* was separated from *A. caudatus* and *A. hypochondriacus* (Figure 2B). The phylogenetic tree presented four major clades for these eight species; one contained the outgroup *A. spinosus*; one the tetraploid *A. dubius*; one was the grain amaranth complex plus *A. blitum*, and one was *A. tricolor* and *A. viridis* (Figure 2C). Based on the genomic components of the structure analysis at K = 8, *A. blitum* and *A. cruentus* shared similar genomic components, and *A. caudatus* was similar with *A. hypochondriacus* (Figure S1). When K = 9 and 10, *A. blitum* and *A. cruentus* were distinguishable from each other; however, *A. caudatus* and *A. hypochondriacus* still shared a large proportion of genetic components until K = 11 (Figure 2D). As the cross-validation of the structure analysis dropped very slowly after K = 11 (Figure S2), we used the Q matrix of K = 11 to cluster the accessions (Table S3). Besides, the *A. tricolor* germplasm was separated into two genetic groups, sub group I consisted of the majority of *A. tricolor*. An interesting observation was that *A. tricolor* sub group I was genetically distant from sub group II, suggesting a strong population structure within this species. In summary, population structure analysis separated eight amaranth species analyzed in this study into nine genetic groups, and two of them belonged to *A. tricolor*.

**Figure 2.**
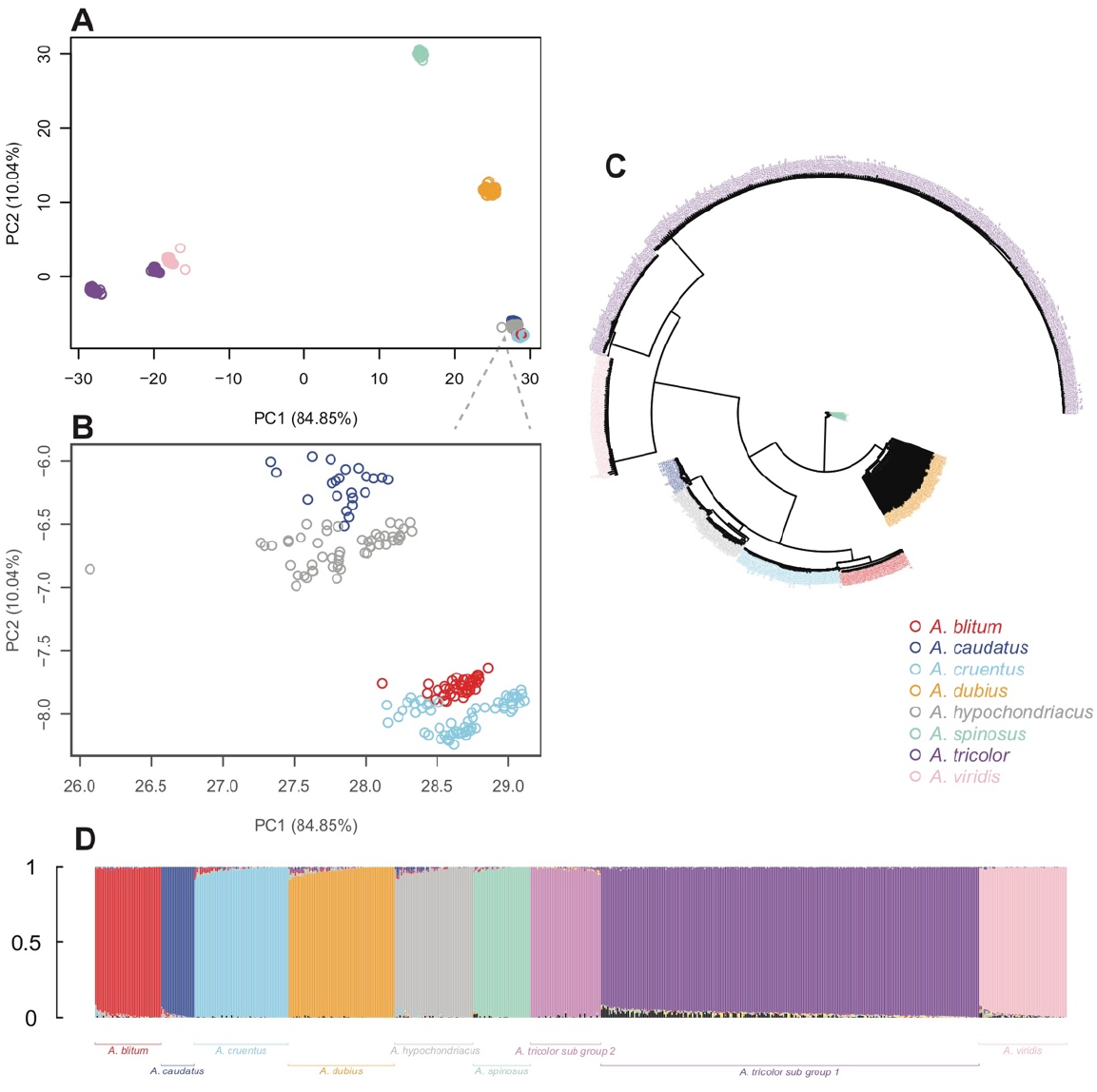
Population structure, phylogenetic tree and structure analysis of the eight *Amaranthus* species. (A)Principal Coordinate Analysis of the eight *Amaranthus* species; (B) Zoom-in into the Principal Coordinate plot of the grain amaranth complex; (C) Phylogenetic tree; (D) Genomic components when K = 11. The color labels are the same across the four plot panels except for an additional light purple in (D) to refer to a subgroup in *A. tricolor*.

Based on the Q matrix of K = 11, 683 accessions showed single-ancestral genomic components (single genetic component > 0.9). Several accessions were found to cluster with a different species than expected based on their original annotation in the passport data. After confirmation by analyzing pictures of the putatively misclassified single-ancestral accessions, we updated the species information of a total of 151 accessions. The accessions with mixed genomic components, which were not assigned into a single genetic group, generally displayed intermediate phenotypes and their species annotation could not be determined with high confidence; therefore, their species record in the passport data was left unchanged. The congruence between species attribution based on morphology and genotype in our collection was about 95% after correction.

### Genetic diversity of *Amaranthus* species

We used the accessions with consistent taxonomy in terms of morphology and genotype to analyze the genetic diversity of the eight major *Amaranthus* species in the WorldVeg collection. Overall, the genetic variance between species was about 86% and the genetic variance within species was about 3%, confirming that *Amaranthus* species are genetically distinct from each other. The observed heterozygosity of *Amaranthus* species was generally low (Table 2), which is reasonable because *Amaranthus* is mostly inbreeding. The observed heterozygosity and the gene diversity within-population (expected heterozygosity) of *A. dubius* was the highest among the *Amaranthus* species (Table 2), resulting from its allotetraploid nature. In grain amaranth complex, *A. caudatus* showed the highest private allele per accession, implying the potential of this germplasm to broaden the genetic diversity of amaranths. The high private alleles per accession of *A. tricolor* and *A. viridis* were corresponding to the relatively distant genetic distance from other *Amaranthus* species (Figure 2A and Table 2).

**Table 2.** The genetic diversity of the main *Amaranthus* species in this collection

| Species | Accession Number | Ho | Hs | PA | Average PA | S | $\pi$ | $\theta$ |
| --- | --- | --- | --- | --- | --- | --- | --- | --- |
| *A. blitum* | 44 | 0.0050 | 0.0497 | 117 | 2.66 | 459 | 0.0038 | 0.0072 |
| *A. caudatus* | 22 | 0.0066 | 0.0588 | 300 | 13.64 | 571 | 0.0125 | 0.0109 |
| *A. cruentus* | 63 | 0.0053 | 0.0474 | 89 | 1.41 | 637 | 0.0049 | 0.0091 |
| *A. dubius* | 71 | 0.2562 | 0.1766 | 1,013 | 14.27 | 6,149 | 0.1682 | 0.0867 |
| *A. hypochondriacus* | 53 | 0.0064 | 0.0557 | 392 | 7.40 | 1,217 | 0.0201 | 0.0183 |
| *A. spinosus* | 38 | 0.0092 | 0.1315 | 337 | 8.87 | 849 | 0.0151 | 0.0141 |
| *A. tricolor* | 300 | 0.0036 | 0.0492 | 3,158 | 10.53 | 4,193 | 0.0568 | 0.0447 |
| *A. viridis* | 59 | 0.0059 | 0.0823 | 1,206 | 20.44 | 1,008 | 0.0087 | 0.0148 |

### Diverged phenotypes between grain and leaf amaranths

The World Vegetable Center genebank applies a standardized documentation of morphological descriptors during the seed regeneration process. Some of these descriptors such as inflorescence and seed morphology reflect important agronomic traits. The measured traits showed no obvious clustering across years in PCA, and nor did the climate parameters (Figure S3), suggesting even in absence of any reference lines it can be assumed that there has been no systematic bias for the phenotypes evaluated across the different years. For each of the categorical traits seed color, inflorescence color, terminal inflorescence shape, petiole pigmentation, leaf shape and pigmentation more than five categories were found (Figure 3), implying high phenotypic diversity of the germplasm set. Besides, some accessions presented more than one categorical phenotype for a single trait, suggesting phenotypic plasticity or heterogeneity in the material. Most quantitative traits showed a skewed distribution over the germplasm (Figure S4), which may reflect different phenotypic distributions among species given that the sample size of leaf amaranth in this collection was larger than the grain amaranth. To investigate the phenotypic divergence among species, we summarized the mean of the quantitative traits of each major *Amaranthus* species in the collection (Table 3). Generally, the grain amaranths (*A. caudatus* and *A. hypochondriacus*) showed early flowering, longer branches and larger inflorescences, but also more seed shattering and smaller grains than the leaf amaranths (*A. blitum*, *A. dubius*, *A. tricolor* and *A. viridis*) (Figure 4). As for dual-purpose amaranth, *A. cruentus* plants in average are higher, have larger leaves and also flower later than grain and leaf amaranths (Figure 4). *A. hypochondriacus* showed similar phenotypes as *A. caudatus* except for its longer branches (Figure S5). *A. blitum* and *A. dubius* presented larger inflorescences than *A. tricolor* and *A. viridis; A. tricolor* showed larger leaves and later flowering than *A. viridis* (Figure S5).

**Figure 3.**
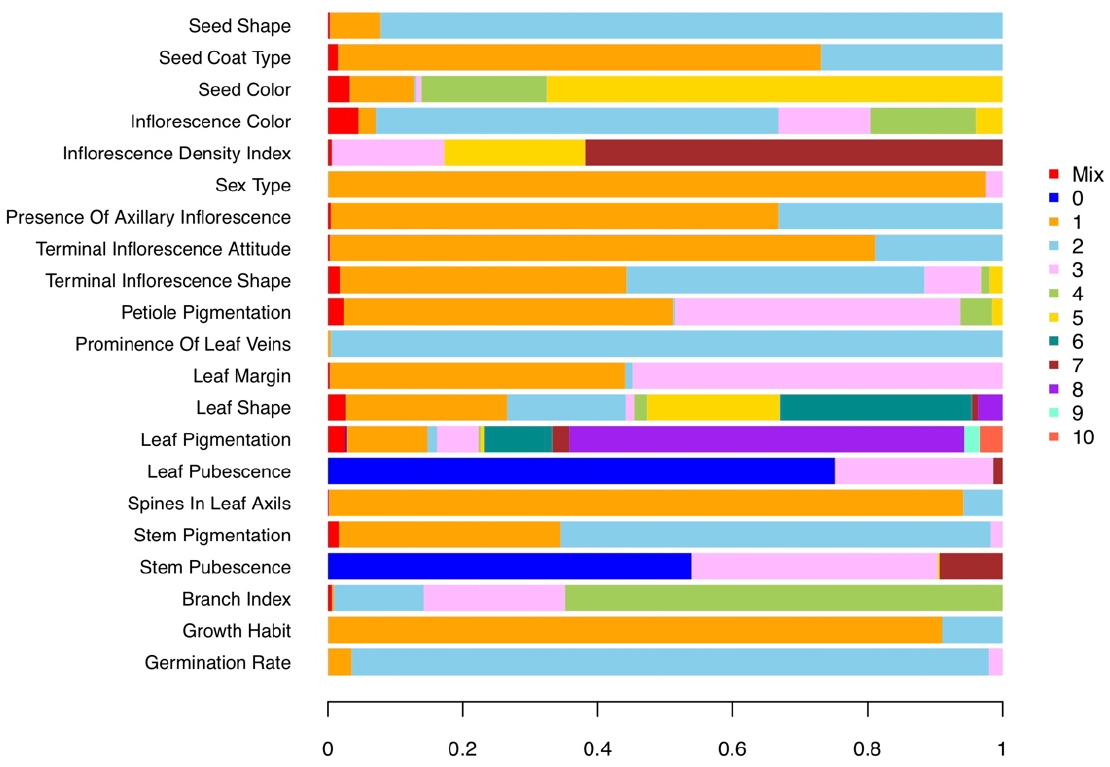
The percentage of each category of the categorical traits. The descriptor codes are Germination rate: 1 = Rapid (< 2 days), 2 = Slow (2-7 days), 3 = Very slow (> 7 days), 4 = Irregular; Growth habit: 1 = Erect, 2 = Prostrate; Branching index: 1 = No branches, 2 = Few branches (all near the base of the stem), 3 = Many branches (all near the base of the stem), 4 = Branches all along the stem; Stem pubescence: 0 = None, 3 = Low, 7 = Conspicuous; Stem pigmentation: 1 = Green, 2 = Purple or pink, 3 = White, M = Mixture; Spines in leaf axils: 1 = Absent, 2 = Present, M = Mixture; Leaf pubescence: 0 = None, 3 = Low, 7 = Conspicuous; Leaf pigmentation: 1 = Entire lamina purple or pink, 2 = Basal area pigmented, 3 = Central spot, 4 = Two stripes (V-shaped), 5 = One stripe (V-shaped), 6 = Margin and vein pigmented, 7 = One pale green or chlorotic stripe on normal green, 8 = Normal green, 9 = Dark green,10 = Other, M = Mixture; Leaf shape: 1 = Lanceolate, 2 = Elliptical, 3 = Cuneate, 4 = Obovate, 5 = Ovatainate, 6 = Rhombic, 7 = Oval, 8 = Other, M = Mixture; Leaf margin: 1 = Entire, 2 = Crenate, 3 = Undulate, 4 = Other, M = Mixture; Prominence of leaf veins: 1 = Smooth, 2 = Rugose (veins prominent); Petiole pigmentation: 1 = Green, 2 = Dark green, 3 = Purple, 4 = Dark purple, 5 = White, M = Mixture; Terminal inflorescence shape: 1 = Spike, 2 = Panicle with short branches, 3 = Panicle with long branches, 4 = Club-shaped at tips, 5 = Other, M = Mixture; Terminal inflorescence attitude: 1 = Erect, 2 = Dropping; Presence of axillary inflorescence: 1 = Absent, 2 = Present; Sex type: 1 = Monoecious, 2 = Dioecious, 3 = Polygamous; Inflorescence density index: 3 = Lax, 5 = Intermediate, 7 = Dense; Inflorescence color: 1 = Yellow, 2 = Green, 3 = Pink, 4 = Red, 5 = Other, M = Mixture; Seed color: 1 = Pale yellow, 2 = Pink, 3 = Red, 4 = Brown, 5 = Black, M = Mixture; Seed coat type: 1 = Translucent, 2 = Opaque, M = Mixture; Seed shape: 1 = Round, 2 = Ellipsoid or ovoid.

**Table 3.** Mean and standard error of eleven quantitative traits of the main eight *Amaranthus* species in this collection. The descriptor codes are Am220: plant height at flowering stage (cm); Am240: mean length of basal lateral branches (cm); Am250: mean length of top lateral branches (cm); Am290: leaf length (cm); Am300: leaf width (cm); Am400: days to flowering (from sowing to 50% with inflorescence); Am410: terminal inflorescence stalk length (cm); Am420: terminal inflorescence laterals length (cm); Am460: length of axillary inflorescence (cm); Am500: seed shattering; 0 = no shattering, 1 = low (<10%), 2 = intermediate (10-50%), 3 = high (>50%); Am540: 1,000 seeds weight (gm).

| Species/Trait Code | Accession Number | Am220 | Am240 | Am250 | Am290 | Am300 | Am400 | Am410 | Am420 | Am460 | Am500 | Am540 |
| --- | --- | --- | --- | --- | --- | --- | --- | --- | --- | --- | --- | --- |
| *A. blitum* | 32 | 159.09±5.70 | 61.48±6.18 | 38.22±4.87 | 17.95±0.52 | 8.88±0.20 | 35.53±1.52 | 29.41±2.17 | 14.93±2.36 | 13.66±0.55 | 3 | 0.19±0.02 |
| *A. caudatus* | 15 | 102.93±8.58 | 9.32±2.52 | 13.71±3.06 | 13.03±1.00 | 7.52±0.54 | 35.00±2.07 | 26.04±2.52 | 9.83±2.41 | 10.14±1.52 | 3 | 0.53±0.09 |
| *A. cruentus* | 58 | 159.41±9.40 | 44.67±4.03 | 18.25±1.60 | 20.62±0.84 | 9.70±0.34 | 63.04±3.08 | 22.10±1.76 | 13.83±0.99 | 9.63±0.30 | 3 | 0.26±0.02 |
| *A. dubius* | 67 | 112.58±6.01 | 53.94±4.57 | 24.05±2.57 | 13.7±0.50 | 10.35±0.48 | 40.64±1.77 | 27.64±2.14 | 18.32±1.49 | 13.35±0.92 | 2 | 0.29±0.03 |
| *A. hypochondriacus* | 63 | 88.52±5.10 | 31.57±3.22 | 24.19±2.62 | 13.86±0.51 | 7.38±0.28 | 33.17±1.36 | 26.76±1.85 | 15.61±1.70 | 16.72±1.29 | 3 | 0.64±0.04 |
| *A. spinosus* | 40 | 72.60±5.33 | 31.02±3.78 | 6.00±1.83 | 8.39±0.33 | 4.48±0.18 | 46.42±2.89 | 8.22±1.08 | 7.36±0.77 | 7.13±0.55 | 2 | 0.22±0.03 |
| *A. tricolor* | 271 | 56.65±2.17 | 21.96±1.23 | 6.95±0.51 | 12.75±0.29 | 8.67±0.21 | 46.29±1.10 | 9.72±0.54 | 6.28±0.44 | 8.10±0.37 | 2 | 0.76±0.02 |
| *A. viridis* | 58 | 47.95±2.45 | 37.01±2.77 | 9.19±0.94 | 6.71±0.16 | 5.21±0.14 | 35.00±1.38 | 8.45±0.55 | 6.37±0.34 | 5.26±0.28 | 2 | 0.32±0.03 |

**Figure 4.**
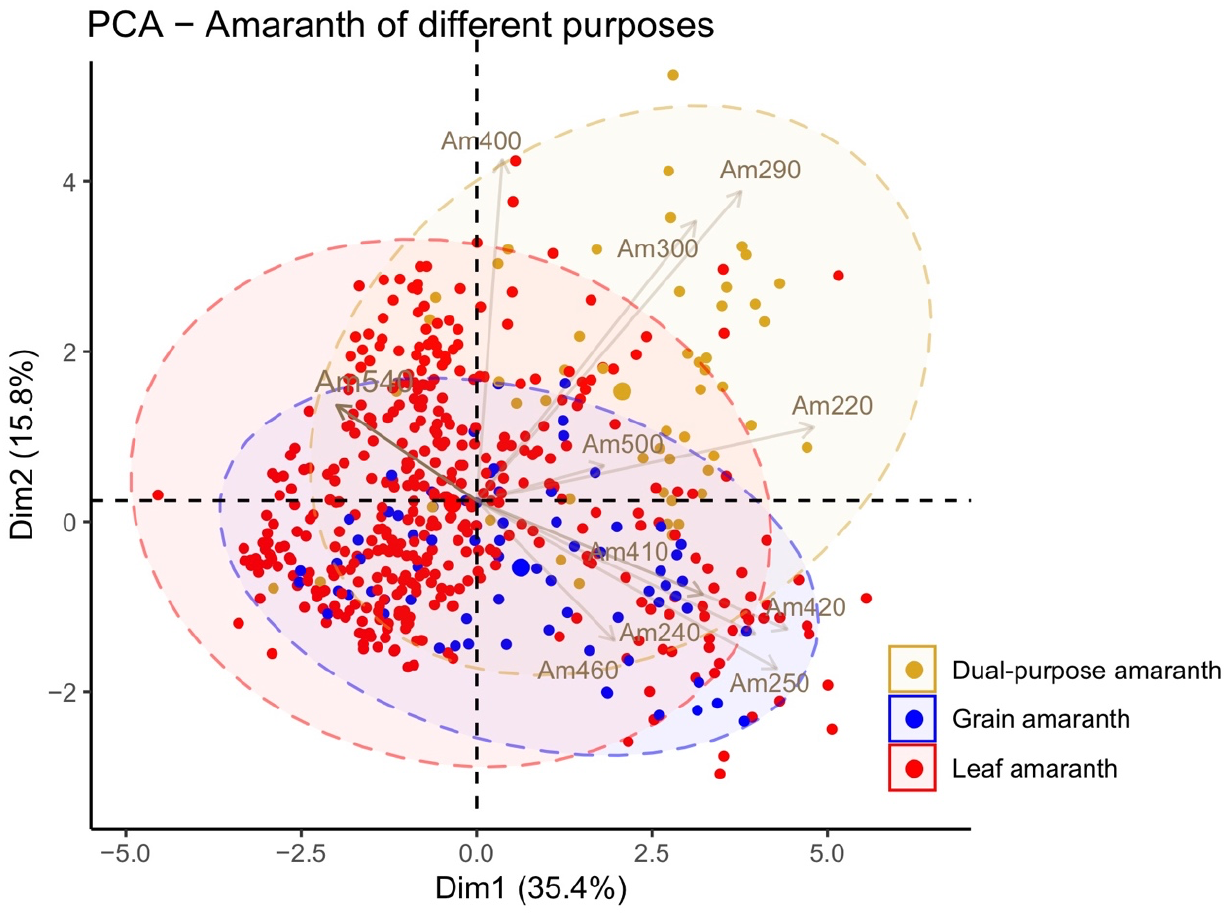
The phenotypic variation of different-purpose amaranth. Grain amaranths contain *A. caudatus* and *A. hypochondriacus;* leaf amaranths contain *A. blitum, A. dubius, A. tricolor* and *A. viridis*. A. cruentus serves as a dual-purpose amaranth. The descriptor codes are Am220: plant height at flowering stage (cm); Am240: mean length of basal lateral branches (cm); Am250: mean length of top lateral branches (cm); Am290: leaf length (cm); Am300: leaf width (cm); Am400: days to flowering (from sowing to 50% with inflorescence); Am410: terminal inflorescence stalk length (cm); Am420: terminal inflorescence laterals length (cm); Am460: length of axillary inflorescence (cm); Am500: seed shattering; 0 = no shattering, 1 = low (<10%), 2 = intermediate (10-50%), 3 = high (>50%); Am540: 1,000 seeds weight (gm).

### Interspecies GWAS of days to flowering

Given that days to flowering is an important trait and has been well documented during germplasm regeneration, we used this trait as a model to investigate the potential of interspecies GWAS in the genus *Amaranthus*. A total of four models, including Phenotype = Genotype, Phenotype = Genotype + PC matrix, Phenotype = Genotype + Q matrix and Phenotype = Genotype + kinship, were used to detected QTL in different panels. First of all, we combined the grain amaranth complex (totally 196 accessions) as a diverse panel because the interspecies hybridization among the grain amaranth complex occurred frequently and naturally. The Q-Q plots of the grain amaranth complex showed a better correction in the model of Phenotype = Genotype + Q matrix (Figure 5A). In this model, we identified 7 significant catalogs, resulting in 12 associated genes on the *A. hypochondriacus* reference genome (Table 4). After blast to Arabidopsis genes, *AH021353-RA* was revealed to be homologous to *AGL20/SOC1* (*AT2G45660*), a gene regulated by *CONSTANS* (*CO*) to promote flowering and by *FLOWERING LOCUS C* (*FLC*) to delay flowering (Lee et al., 2000; Yoo et al., 2005; Lee and Lee, 2010). The second panel contained all the *A. tricolor* accessions (307 accessions). However, none of any model showed a better correction (Figure 5B), suggesting the strong population structure in *A. tricolor* may lead to a serious false positive result. Given that the sub group I of *A. tricolor* contained a total of 253 accessions, we used this panel to perform another GWAS, resulting in a better correction in the models of Phenotype = Genotype + PC matrix and Phenotype = Genotype + Q matrix (Figure 5C). Combining the results of these two models, a total of 28 *A. hypochondriacus* genes were associated with 11 significant catalogs (Table 5). We identified six candidate genes that were homologous to Arabidopsis genes regulating flowering time. *AH018224* was homologous to *low-beta-amylase 1* (*lba1; AT5G47010*) which showed early flowering in the Arabidopsis mutant (Yoine et al., 2006). *AH021320, AH021553* and *AH001353* were homologous to *brassinosteroid*-*insentitive 1*(*bri1; AT4G39400*), an enhancer of *FLC*, consequently leading to late flowering (Domagalska et al., 2007). *AH021139* was homologous to *sgs1* (*AT3G10490*), which encoded a transcription factor of *NAC52* and regulated flowering time through DNA methylation (Butel et al., 2017). *AH001354* was homologous to *FLOWERING TIME CONTROL LOCUS A* (*FCA; AT4G16280*); a *fca1* mutant was revealed late flowering in Arabidopsis (Macknight et al., 1997). Together with the GWAS result of the grain amaranth complex, the functions of these seven candidate genes could be confirmed through further molecular experiments.

**Figure 5.**
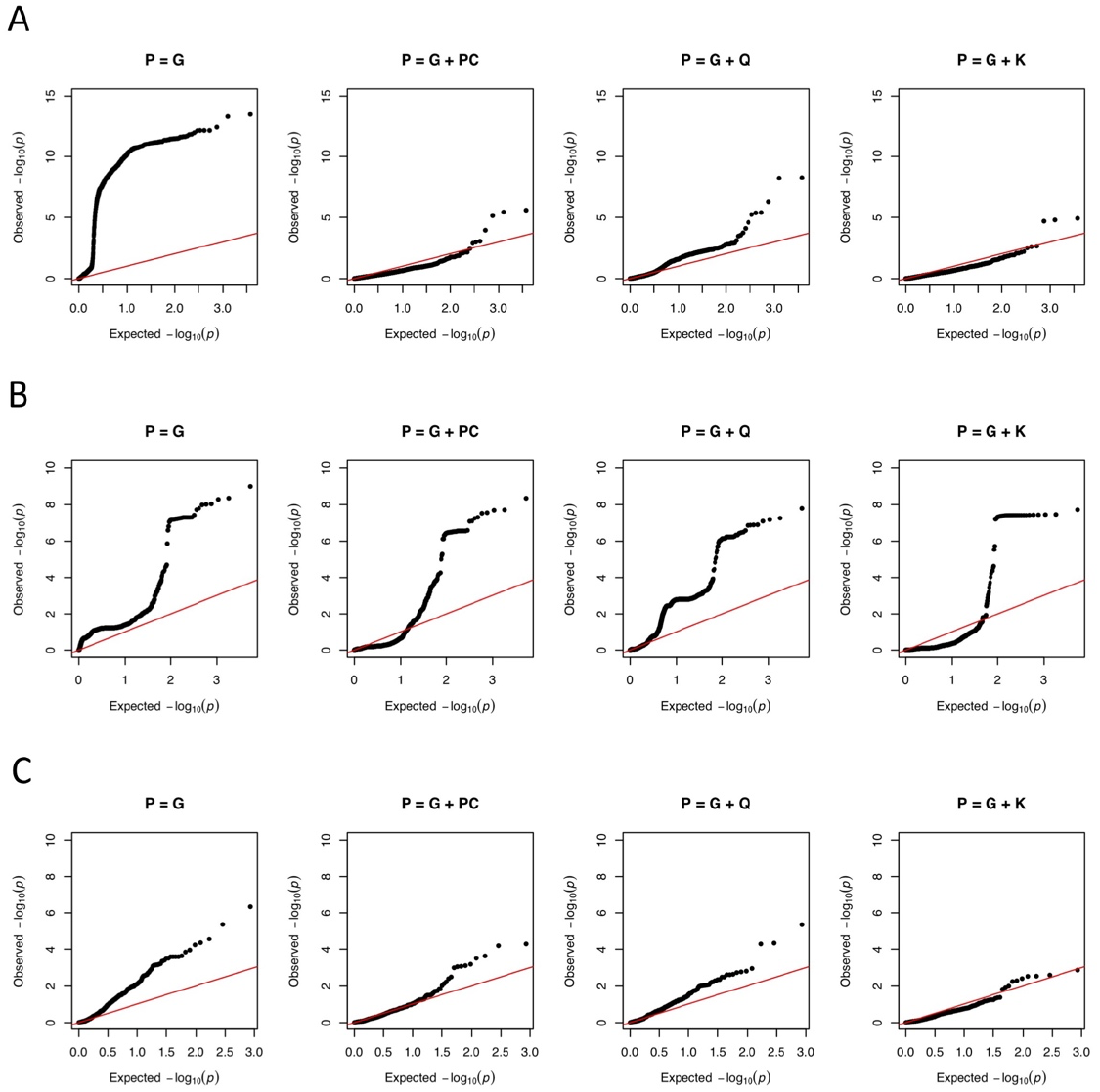
The Q-Q plots of different GWAS panels: A) grain amaranth complex, including *A. caudatus*, *A. cruentus* and *A. hypochondriacus*; B) *A. tricolor*; C) the sub group I of *A. tricolor*.

**Table 4.**
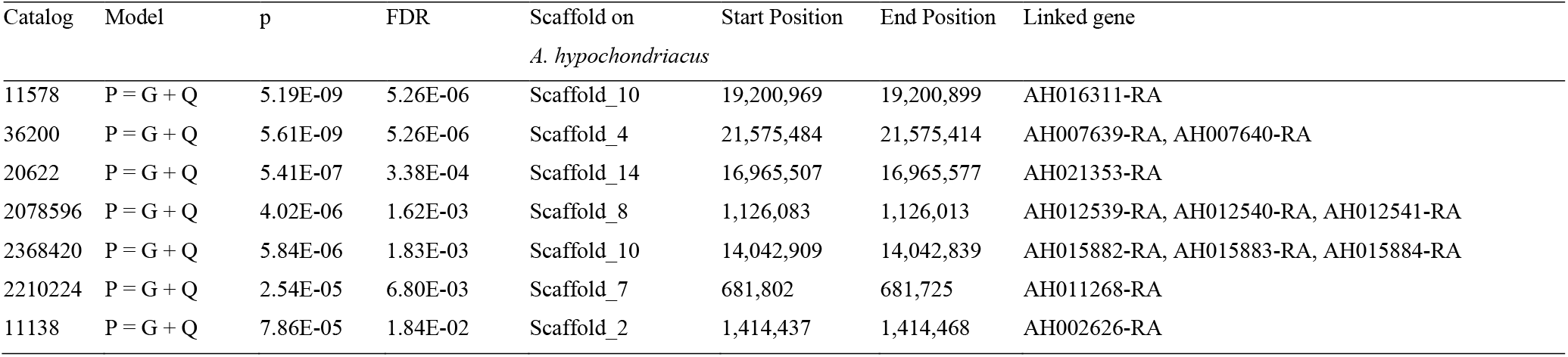
Candidate SNPs of interspecific GWAS (grain amaranth complex) for days to flowering

**Table 5.**
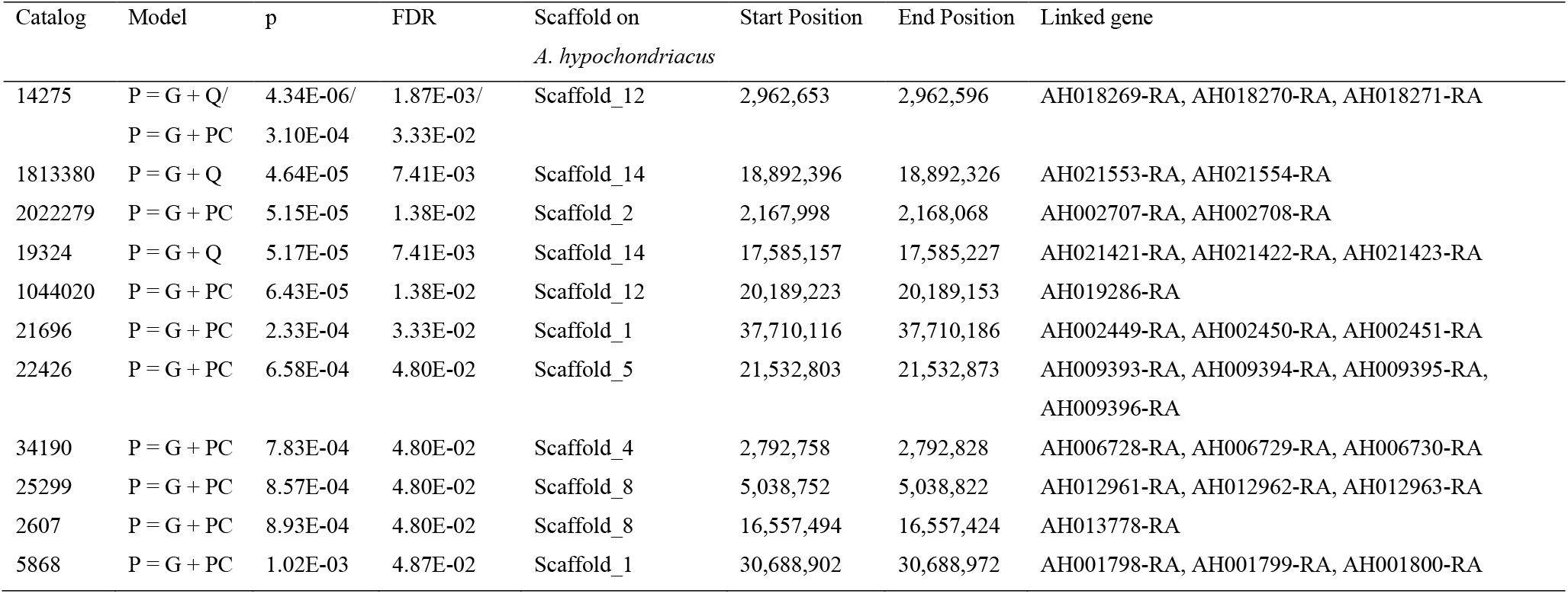
Candidate SNPs of GWAS based on the sub group I of *A. tricolor* for days to flowering

## Discussion

Genebank collections around the world hold large numbers of amaranth accessions. The largest collection is hosted by the ICAR-National Bureau of Plant Genetic Resources, India, with a base collection of 5,804 accessions (Pandey et al., 2016). The United States Department of Agriculture, North Central Regional Plant Introduction Station holds 3,342 accessions, Embrapa Brasil 2,495 accessions, and the Institute for Agrobotany in Tapioszele, Hungary, holds 787 accessions (Genesys, 2017). The World Vegetable Center currently holds 1,062 accessions. Access to correct passport data and genotypic and phenotypic information for genebank accessions likely increases the use of this material for research, breeding and direct cultivation (Anglin et al., 2018). For many germplasm collections held in genebanks, especially for less utilized crops such as amaranth, only very basic passport data are available, which may be incomplete or contain errors, while characterization data are completely missing. Documenting the results of germplasm characterization during seed regeneration is an efficient means to generate information on plant morphology and of basic agronomic characteristics of genebank accessions (Philipp et al., 2018). Likewise, genotypic characterization of germplasm collections provides insights into the diversity and population structure of the collection, helps to identify duplicates as well as collection gaps in the genebank collections, and, if genotyping is performed across genebanks, gives a global overview of the *ex-situ* conservation of a crop (Mascher et al., 2019). Ideally, the germplasm collections also should be phenotyped, but this task is labor intensive, costly and complex and requires specific strategies to be successfully applied on genebank collections (Nguyen and Norton, 2020).

In the current study we genotyped the WorldVeg *Amaranthus* genebank collection, which comprised almost 800 accessions at the time the study was initiated, and used the genotype data together with historical characterization data to elucidate the diversity and population structure of the collection, map candidate genes for time to flowering and revise erroneous passport data. Previous studies have shown that the taxonomic classification of Amaranthus can be confusing, as it relies on a few characteristics to distinguish species from each other (Sauer, 1950, 1967). El-Ghamery et al. (2017) evaluated 38 leaf anatomical characteristics in the genus *Amaranthus* and proposed eight identification keys (El-Ghamery et al., 2017). However, species identification based on these anatomical features requires specialized skills to prepare the samples. Genebank curators in multi-species genebanks cannot be taxonomic experts for all the crops they manage, so accessions are often only identified at the genus level, or the species annotation may be erroneous. However, for using germplasm in research and breeding, correct species information is key. Therefore, a genotype-based identification system for difficult to determine species would make the species annotation in passport data more reliable and thereby improve the utilization of the conserved germplasm. In this study, we used genetic clustering as supportive evidence to verify species attributions, given that genetic variance between *Amaranthus* species is greater than within species (Wu and Blair, 2017). A 15K SNP dataset together with pictures of the accessions was used to re-examine the species information of the passport data and for single ancestral accessions genotypes proved to be highly practical for correct species annotation. But still, for genetically admixed accessions phenotypic and genotypic species attribution did not match and the species attribution remained unresolved.

*A. dubius* showed the highest observed heterozygosity and gene diversity within-population. This is very likely due to the tetraploid of this species (Grant, 1959), where sequencing reads from different sub-genomes are assembled in the same catalogs in the *de novo* SNP calling pipeline. Previous studies revealed similar extents of gene diversity within-population among grain amaranth complex, while *A. cruentus* had more private alleles than other grain amaranths (Kietlinski et al., 2014; Wu and Blair, 2017). However, in this study, the grain amaranth complex indeed showed similar gene diversity within-population with *A. cruentus* having the fewest private alleles, implying the close kinship among *A. cruentus* accessions in this collection. Besides, it should be noted that the number of private alleles discovered through the ddRAD sequencing approach might be underestimated because of the heterogeneity of restriction enzyme recognition sites (Cariou et al., 2016; Davey et al., 2013). It has been shown that *A. blitum, A. dubius* and *A. tricolor* have larger genome sizes (about 700 Mb) than the other amaranth species (about 500 Mb) (Stetter and Schmid, 2017), and the total sample size in GBS of large-genome species is larger than in small-genome species. Under these circumstances, the private alleles of a small-genome species are very likely to be filtered out due to high missing rate, resulting in the underestimation of their genetic diversity. Nevertheless, the large number of private alleles found in *A. caudatus* could provide new genetic variation for amaranth breeding via interspecies hybridization.

To our knowledge, rare studies addressed the genetic diversity of leafy amaranths. Nguyen et al. (2019) identified two genetic groups for *A. tricolor*, one comprising the accessions from America and one for accessions from South Asia and other regions (Nguyen et al., 2019). Also, the WorldVeg *A. tricolor* collection fell into two genetic groups; the accessions from Bangladesh clustered into one individual group and the other contained the rest of the accessions from South and Southeast Asia. Considering that the two clades of *A. tricolor* are diverged from VI048200, an accession from Bangladesh, VI048200 may be the ancestor of local varieties in South and Southeast Asia (Figure 2D). Leaf amaranths, especially *A. tricolor*, are popular vegetables in Asia. *A. tricolor* can be cultivated in the open field, in the greenhouse, or in plant factories, and can be sold as up-rooted small plants, or as a microgreen (Ebert et al., 2014). Due to its short rotation time, it easily fits into crop calendars and is a good option when only a short growing window is available. The versatility of the crop in adapting to various production systems and its high nutrient content warrants more investigation in genetic and phenotypic characteristics of the germplasm to develop improved varieties for a broad range of utilizations.

Phenotype-driven utilization of a crop genus or species for different purposes is common in plants. For example, the genus *Brassica* comprises vegetable crops, such as *Brassica oleracea*, or oil crops, such as *Brassica rapa*, which also includes vegetable crops such as turnip and bok choy (Cheng et al., 2016; Lin et al., 2014; Qiao et al., 2020). In this study, we propose that the utilization of amaranths for different purposes may result from the phenotypic divergence among different species, given that amaranth is an ancient crop and has been consumed in various ways in different regions. Amaranth likely has been consumed by early humans and amaranth seed was found in early neolithic settlements in Europe, together with seeds of cereals, legumes and fruits (Marinova et al., 2002). The oldest findings of domesticated grain amaranth in the Americas were detected in the Coxcatlán cave in Mexico dating from 2,000 BCE (Schultze-Motel, 1972). Farmers in Mesoamerica may have preferred to harvest grains rather than leaves from the early flowering amaranths with larger inflorescences, which seem to be an essential element of grain amaranths and may be able to compensate the unattractive lower seed weight (Figure 4 and Table 3). In contrast, the small inflorescences of leaf amaranths are unlikely attractive to farmers pursuing high grain yield. However, late flowering means a later shift from vegetative to the reproduction stage; as a result, nutrients are allocated to leaves over a longer time and farmers can harvest more leaves. It is not possible to distinguish leaf and grain amaranths purely by species - there are grain amaranth species that are used for vegetable production. But phenotypic-driven preference could explain why *A. cruentus* has been chosen as a dual-purpose amaranth given that generally it is late flowering but has large leaves; hence, farmers could harvest leaves multiple times during the longer vegetative stage, which also provides recovery time to enrich grain yield during the cultivation period (Dinssa et al., 2018; Hoidal et al., 2019). Moreover, the phenotypic variation also points out a direction of amaranth breeding (Figure 4). For example, grain amaranths show large variance in seed shattering, suggesting shattering may be improvable (Brenner and Hauptli, 1990). Reduction of seed shattering is very important, as plants need to dry in the field before harvest, a process that can take weeks. If the genotype has the tendency to shatter seed, the yield loss can be very high (Fitterer et al. 1996). Dual-purpose amaranth shows great variance in most of these traits, suggesting that a desired combination of large leaf, late flowering, long inflorescence and low seed shattering could be achieved.

Genebank genomics is an effective approach to mine genetic resources for trait diversity, also in orphan crops. However, unlike major crops for which large numbers of accessions and data have been accumulated, only a relatively small number of accessions are available in genebanks for orphan crops such as amaranth. The amaranth accessions held by the WorldVeg genebank comprise materials of 13 different species, and the number of accessions available of a single species is too small for a GWAS panel. However, interspecific hybridization among *Amaranthus* species occurs naturally and artificial hybridization could introduce new genetic variation into different species; we therefore proposed the interspecies GWAS of grain amaranth complex to identify QTLs regulating attractive agronomic traits in this study. Combining various species in a GWAS panel makes SNP calling using reference sequences difficult because reads from genetically distant species may not be mapped back onto the reference genome, leaving many unmapped reads. The present study applied *de novo* SNP calling to achieve SNPs across several species of the *Amaranthus* genus, keeping more mapped reads and avoiding the risk of losing SNPs located in genomic regions that are absent in the reference sequence (Table 1). Besides, although the reference genome-based SNP calling could estimate LD and would inform about the QTL interval, the detection of associations would be strongly reduced by the structural variation among different species. Considering the LD in a diverse panel would result from the physical linkage and also from the consequence of natural selection of functionally related genes, the estimation of LD between markers produced by ddRAD sequencing that depends on the conservation of restriction enzyme cutting sites would be less reliable. Another issue while combining multiple species into a GWAS panel is the risk of introducing strong population structure, which could result in false positive associations. Here, we mitigated the risk of obtaining false positive associations by introducing structure and kinship as cofactors into the GWAS models, resulting in one candidate genes out of seven significant catalogs in grain amaranths panel, and six candidate genes out of eleven significant catalogs in the sub group I of *A. tricolor*, respectively. QTL mapping in artificial populations produced from divergent species and GWAS in a single species could provide more evidence to confirm our candidate genes from the grain amaranth complex GWAS. These candidate genes could be also supported by further functional validation of the candidate genes in *Amaranthus* species.

## Conclusion

The genus *Amaranthus* contains species that are suitable as grain crop, vegetable crop or dual use crop, combining the use of vegetable and grain harvest. In total 731 accessions of the WorldVeg amaranth collection comprising 777 accessions at the time of the study were genotyped and the genotypic data were used to analyze the diversity and population structure in the collection. Together with pictures documenting the plant morphology, the population structure data were successfully used for correcting the species annotation of single ancestry accessions of the collection. Two genome-wide association study using different panels revealed a total of seven candidate genes associated with the homologous genes regulating flowering time in Arabidopsis. The genotyped and partly phenotypically characterized WorldVeg amaranth collection is available to users for research breeding and training, as well as for direct cultivation at https://avrdc.org/seed/unimproved-germplasm/ and is distributed through the standard material transfer agreement used by the International Treaty on Plant Genetic Resources for Food and Agriculture.

## Acknowledgement

We thank Dr. Li-Yu Liu in the Department of Agronomy, National Taiwan University for her advice on the statistics for this study and the WorldVeg biotechnology team for DNA extraction, genebank team for providing the data of decades of phenotypic evaluation, and the communication team for image editing. This work received financial support from the German Federal Ministry for Economic Cooperation and Development (BMZ) commissioned by the Deutsche Gesellschaft für Internationale Zusammenarbeit (GIZ) Fund for International Agricultural Research (FIA), grant number: 81219429. Additional support was provided by long-term strategic donors to the World Vegetable Center: Taiwan, UK aid from the UK government, United States Agency for International Development (USAID), Australian Centre for International Agricultural Research (ACIAR), Germany, Thailand, Philippines, Korea, and Japan.

## Supplementary figures

Figure S1 Structure analysis of K = 8, 9 and 10. The accessions are in the same order as in the structure plot of K = 11 (Figure 2D).

Figure S2 Cross-validation of the structure analysis.

Figure S3 Principal Component Analysis of (A) the quantitative traits and (B) the climatic variables across the regeneration years, respectively.

Figure S4 Distributions of the quantitative traits in the germplasm panel.

Figure S5 Principal Component Analysis of the quantitative traits of (A) the grain amaranths and (B) the leaf amaranths, respectively. The trait codes refer to the standard descriptors (http://seed.worldveg.org/download).

## Supplementary tables

Table S1 Species, regeneration year and phenotype evaluations of the WorldVeg amaranth collection.

Table S2 Climatic records in Shanhua, Taiwan during the amaranth regeneration in 1995 to 2019.

Table S3 Sequencing information and genetic components of the amaranth accessions of the WorldVeg genebank collection.

