## Supplementary Figures for "*De novo* SNP calling reveals the candidate genes regulating days to flowering through interspecies GWAS of *Amaranthus* genus"

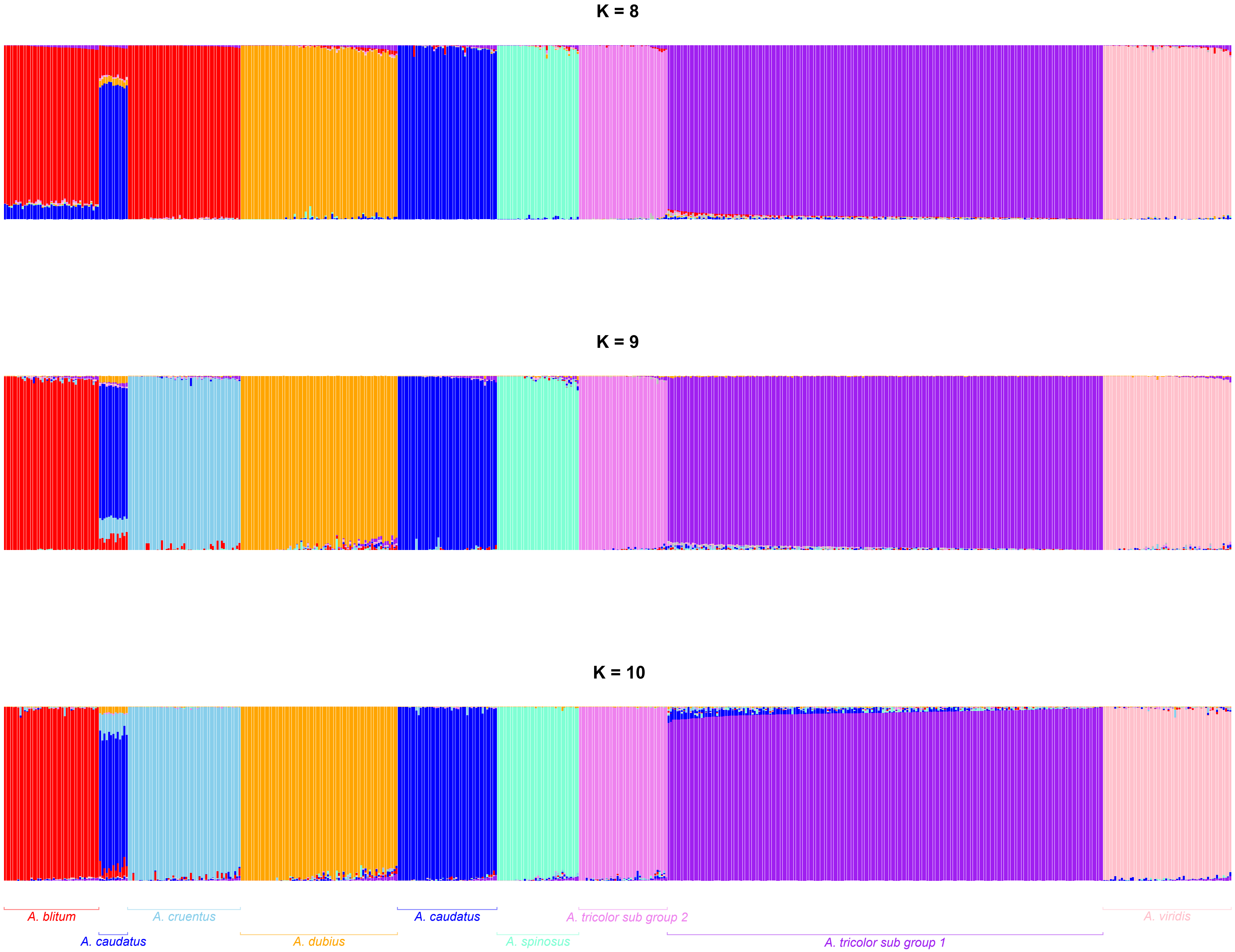


Supplementary Figure 1. Structure analysis of K = 8, 9, and 10. The accessions are in the same order as in the structure plot of K = 11 (Figure 4D).

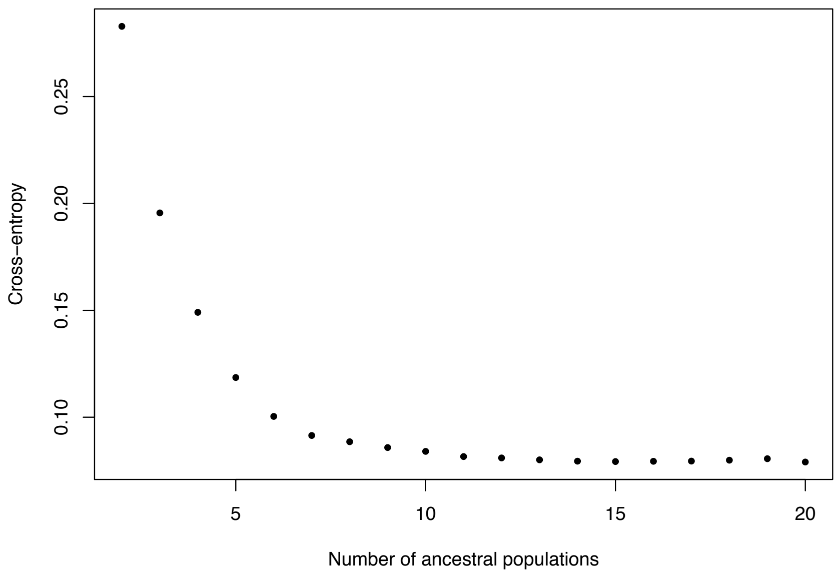


Supplementary Figure 2. Cross-validation of the structure analysis.

(A) PCA of morphological traits in different years

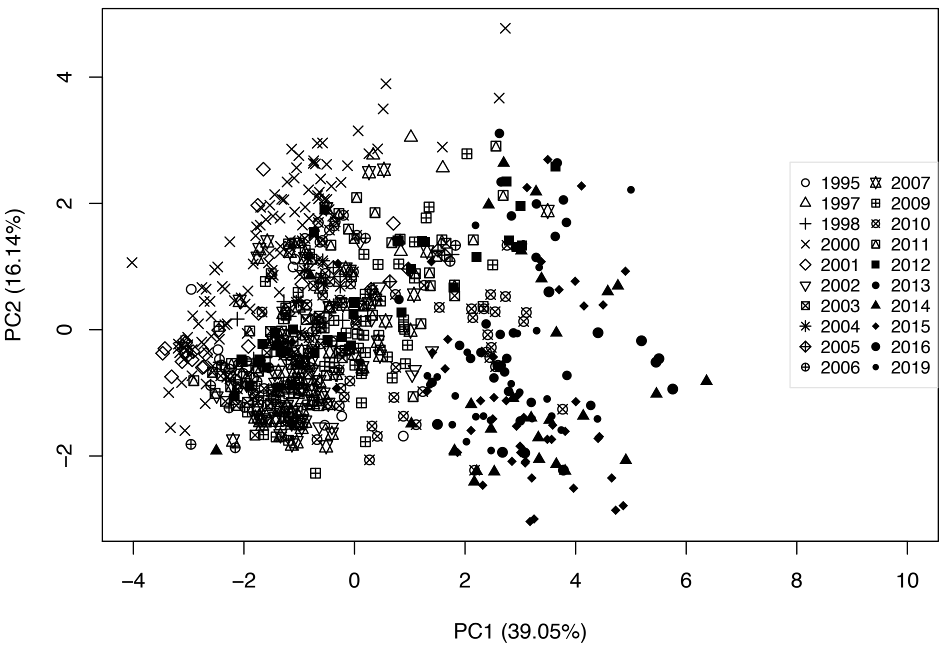


(B) PCA of climatic data in different years

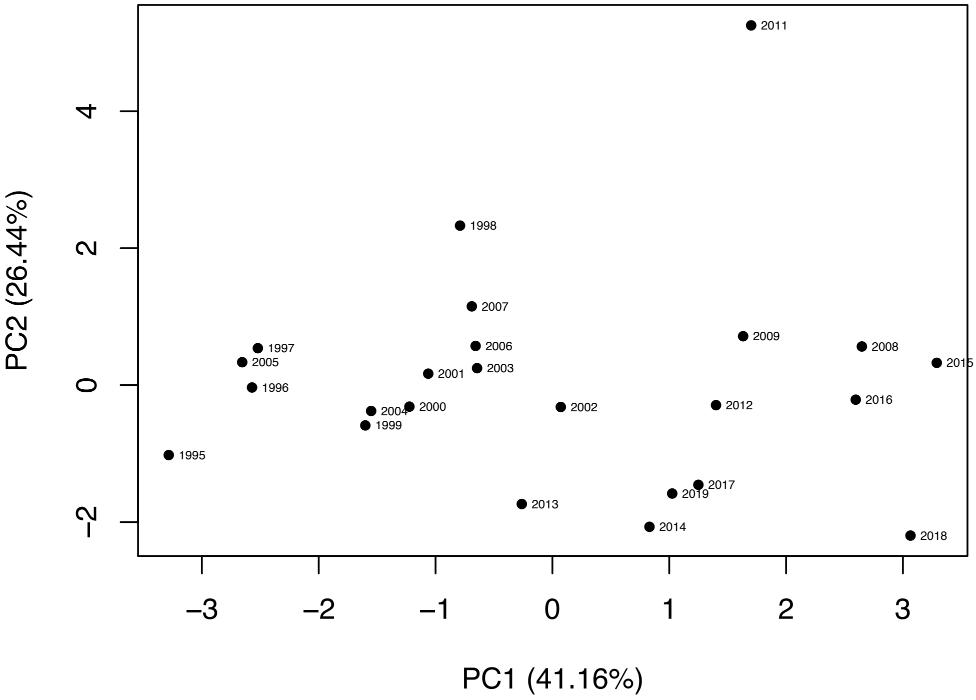


Supplementary Figure 3. Principal Component Analysis of (A) the quantitative traits and (B) the climatic variables across the regeneration years, respectively.

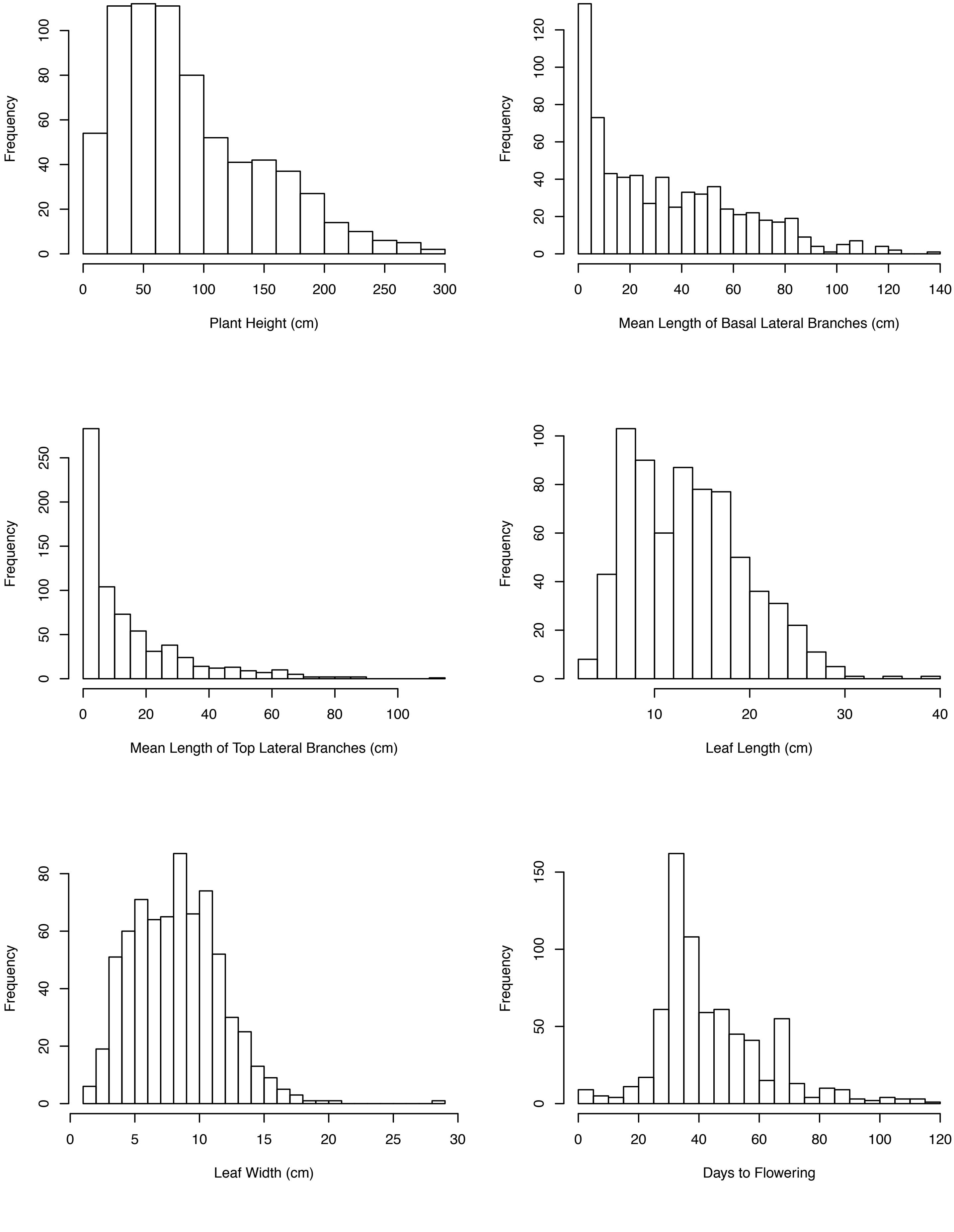

Frequency
Plant Height (cm)
Frequency
Mean Length of Basal Lateral Branches (cm)
Frequency
Mean Length of Top Lateral Branches (cm)
Frequency
Leaf Length (cm)
Frequency
Leaf Width (cm)
Frequency
Days to Flowering

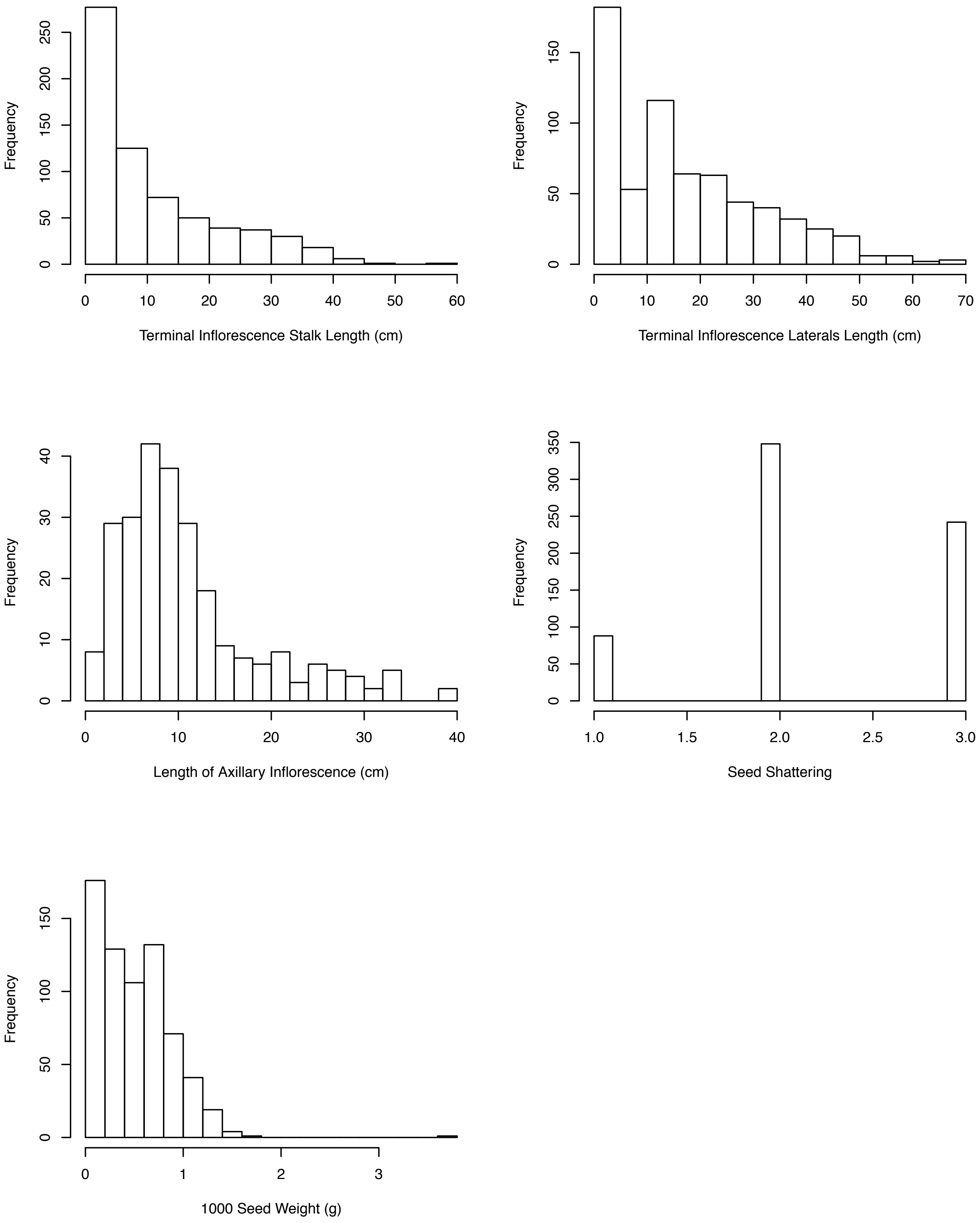


Supplementary Figure 4. Distributions of the quantitative traits in the germplasm panel.

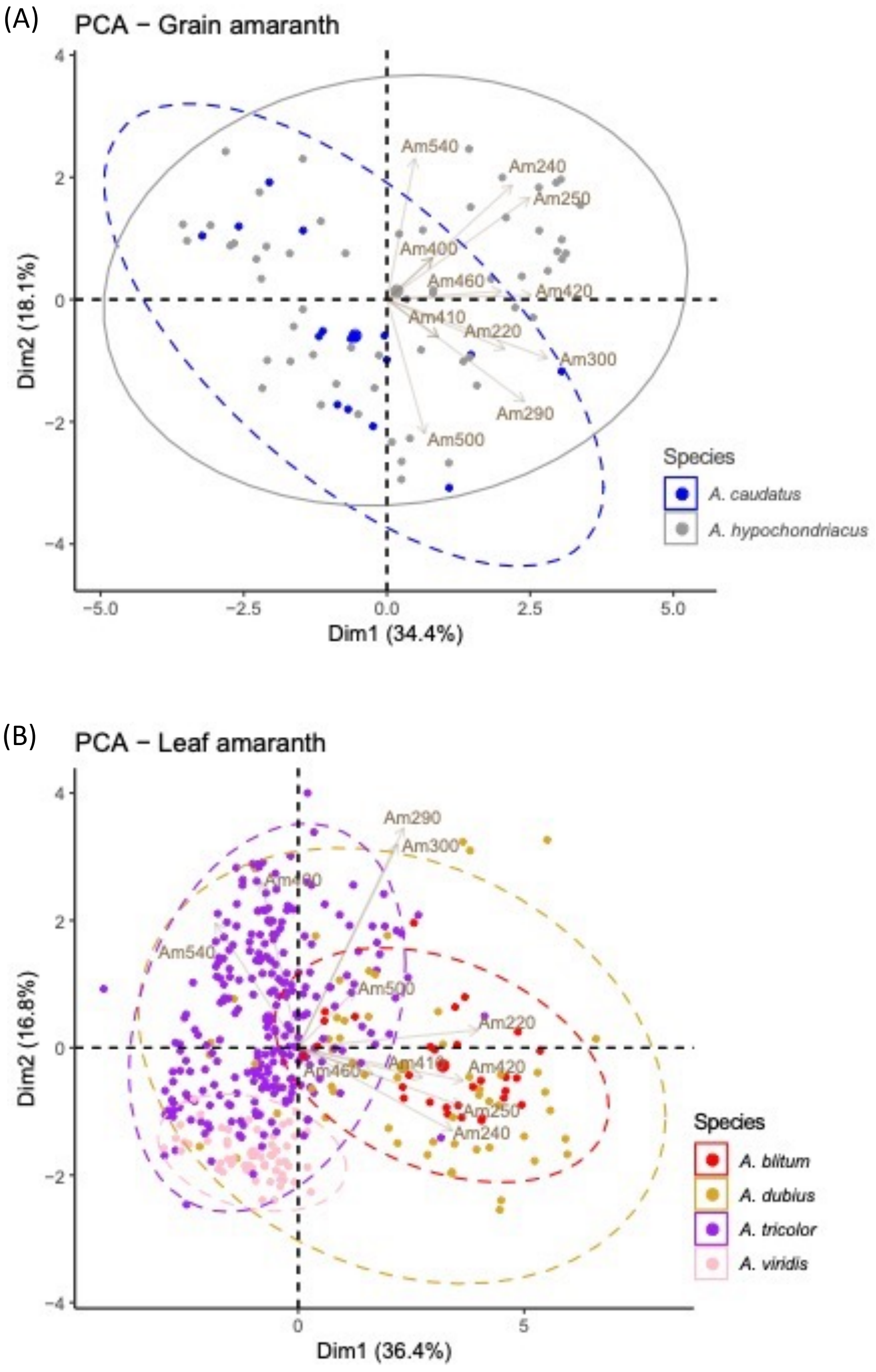


Supplementary Figure 5. Principal Component Analysis of the quantitative traits of (A) the grain amaranths and (B) the leaf amaranths, respectively. The trait codes refer to the descriptor.
